## Supplemental File 1 for "Downregulation of dystrophin expression occurs across diverse tumors, correlates with the age of onset, staging and reduced survival of patients"

**Supplementary Figures**

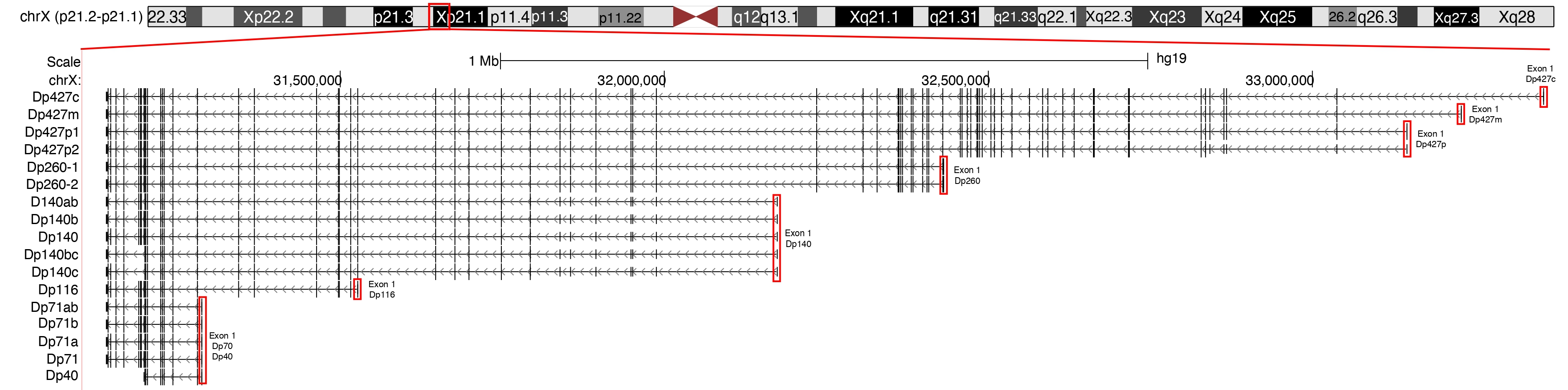

Supplementary Figure 1. Chromosomal localization and transcripts encoded by the *DMD* gene. The vertical dashes indicate individual exons. The location of the first exons is indicated by the red boxes. The full-length dystrophins (Dp427) are encoded by transcripts consisting of 79 exons, with the M, C and P isoforms having unique N-termini encoded by specific first exons. Multiple splice variants have been indicated on the left.

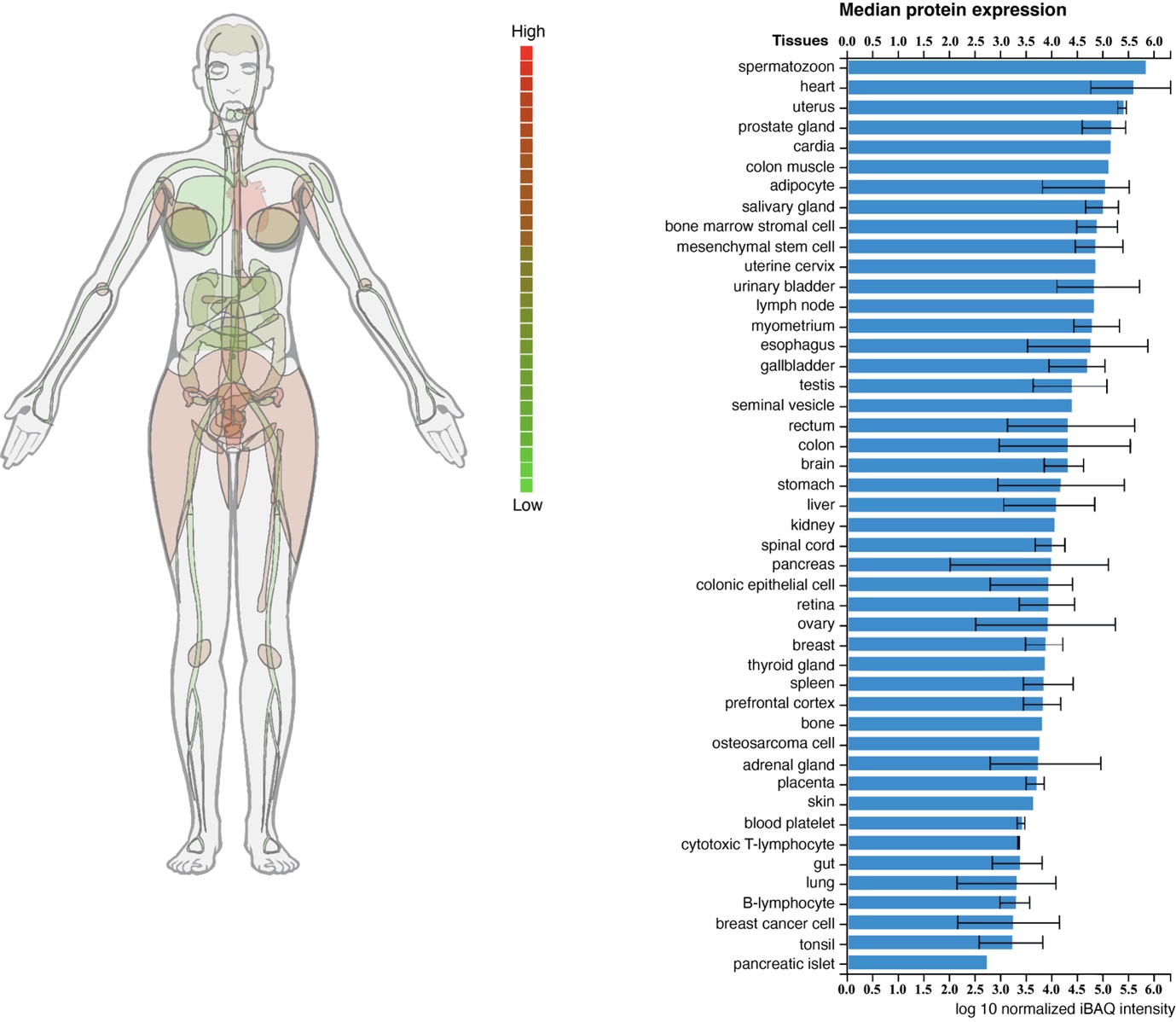

Supplementary Figure 2. Full-length dystrophin protein expression in normal tissues and cells (Proteomics DB).

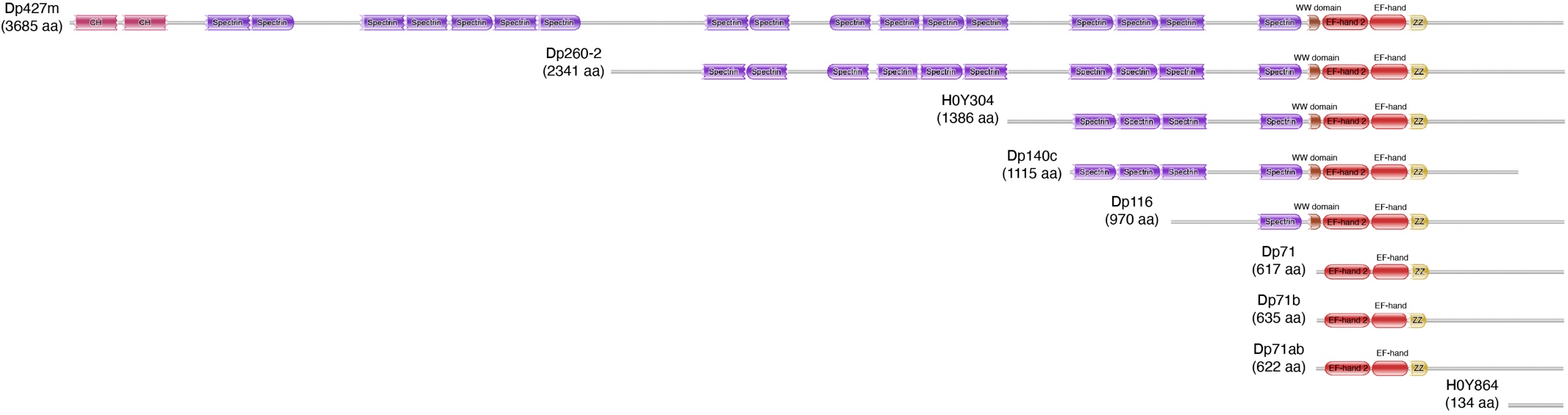

Supplementary Figure 3. The predicted domain structures encoded by *DMD* gene transcripts used in the hierarchical clustering analysis. The figure was obtained using the HMMER web server.

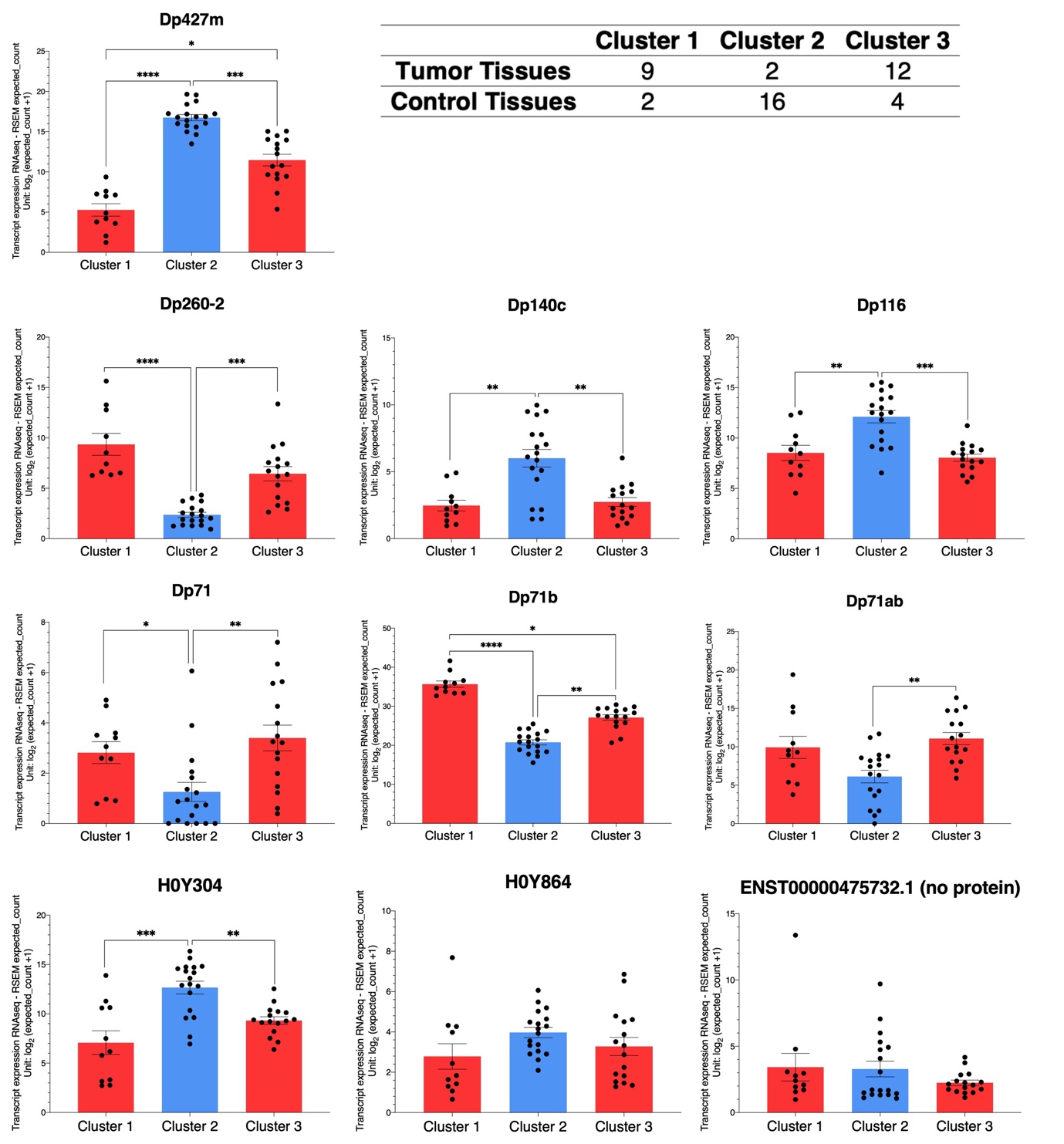

Supplementary Figure 4. Expression level of the top ten highly expressed *DMD* gene transcripts in three of the identified clusters. Expression data is represented as mean ± SEM (**p* <0.05, ***p* <0.01, ****p* < 0.001, *****p* < 0.0001).

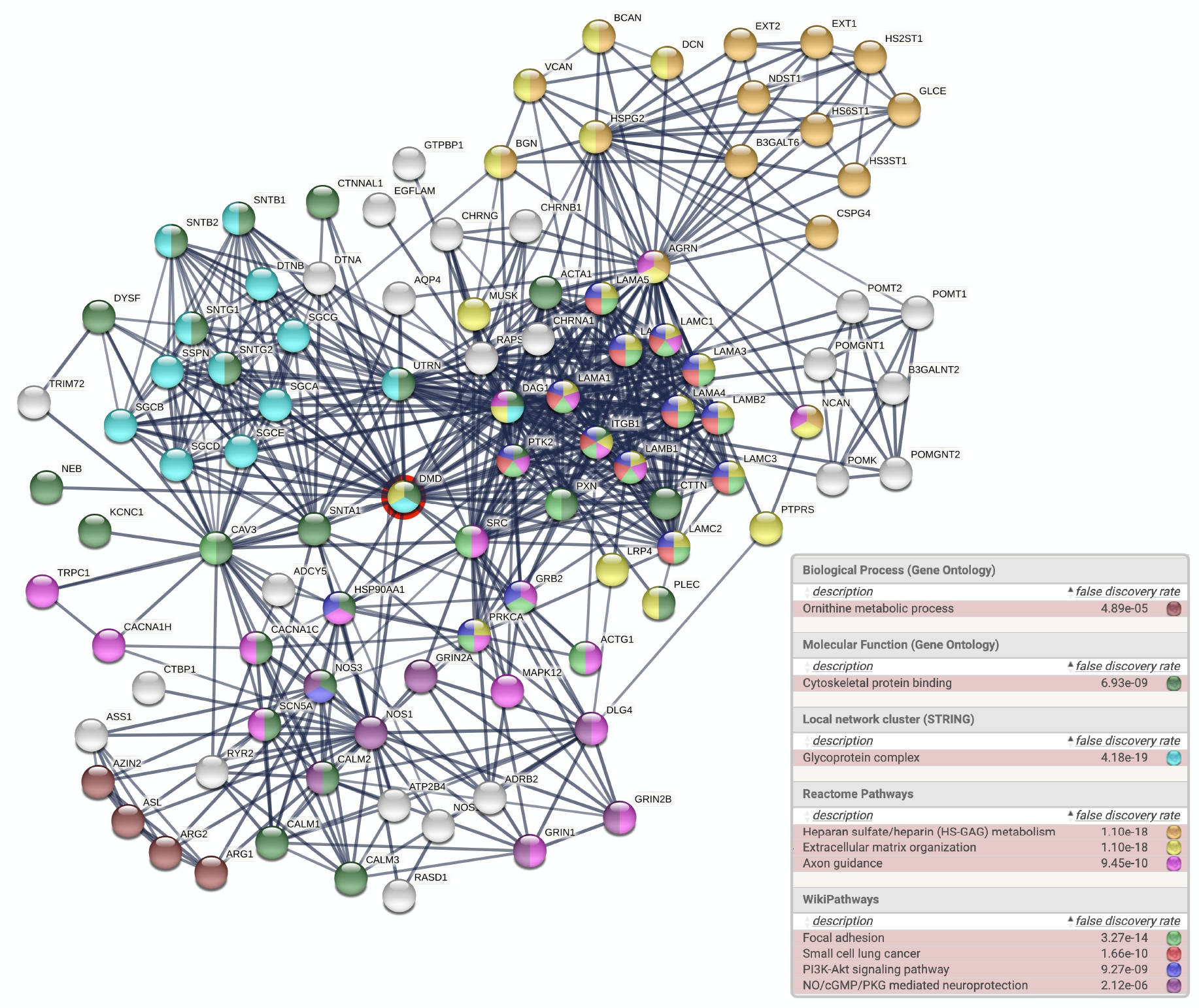

Supplementary Figure 5. STRING interaction diagram of dystrophin protein-protein interaction (PPI) network and the enriched functional terms according to "Reactome Pathways" and "WikiPathways". Each node depicts a gene, edges between nodes indicate interactions between the corresponding protein products.

**Supplementary Tables**

Supplementary Table 1. *DMD* gene expression in 25 TCGA primary tumors and corresponding normal tissues adjacent to the tumors (NATs) from the TCGA or healthy tissues form the GTEx database. The *p*-values were calculated using the UCSC Xena Browser using a two-tailed Welch’s t test and adjusted for multiple testing using the Bonferroni correction. P-values were multiplied by the total numbers of tests (n= 25) and compared with the overall level of α= 0.05. Statistically non-significant comparisons are highlighted in grey.

| **Comparisons with decreased expression of the *DMD* gene in tumor samples** | | | | |
| --- | --- | --- | --- | --- |
| Pairwise comparisons (Number of samples) | | | *DMD* expression | |
|  |  |  | LogFC | *p*-value |
| 1 | Breast invasive carcinoma (1,092) | Normal adjacent tissue (113) | -3.7 | 1.47e^-114^ |
| 2 | Bladder urothelial carcinoma (407) | Normal adjacent tissue (19) | -3.5 | 2.42e^-05^ |
| 3 | Uterine corpus endometrioid carcinoma (180) | Normal adjacent tissue (23) | -3.1 | 1.47e^-14^ |
| 4 | Cholangiocarcinoma (36) | Normal adjacent tissue (9) | -3.1 | 2.69e^-14^ |
| 5 | Cervical and endocervical cancer (304) | GTEx healthy cervix tissue (10) | -2.9 | 5.57e^-5^ |
| 6 | Pancreatic adenocarcinoma (178) | GTEx healthy pancreas tissue (165) | -2.5 | 2.33e^-74^ |
| 7 | Ovarian serous cystadenocarcinoma (419) | GTEx healthy ovary tissue (88) | -2.4 | 8.78e^-84^ |
| 8 | Uterine carcinosarcoma (57) | GTEx healthy uterus tissue (78) | -2.4 | 1.47e^-14^ |
| 9 | Rectal adenocarcinoma (92) | Normal adjacent tissue (10) | -2.4 | 1.04e^-2^ |
| 10 | Colon adenocarcinoma (288) | Normal adjacent tissue (41) | -1.8 | 1.08e^-10^ |
| 11 | Stomach adenocarcinoma (414) | Normal adjacent tissue (36) | -1.7 | 3.42e^-4^ |
| 12 | Prostate adenocarcinoma (495) | Normal adjacent tissue (52) | -1.6 | 4.29e^-14^ |
| 13 | Lung squamous cell carcinoma (498) | Normal adjacent tissue (50) | -1.6 | 7.57e^-48^ |
| 14 | Esophageal carcinoma (181) | GTEx healthy esophagus tissue (651) | -1.6 | 2.78e^-26^ |
| 15 | Skin cutaneous melanoma (102) | GTEx healthy skin tissue (557) | -1.5 | 5.56e^-17^ |
| 16 | Liver hepatocellular carcinoma (369) | Normal adjacent tissue (50) | -1.5 | 1.79e^-36^ |
| 17 | Head and neck squamous cell carcinoma (518) | Normal adjacent tissue (44) | -1.5 | 1.69e^-4^ |
| 18 | Lung adenocarcinoma (513) | Normal adjacent tissue (59) | -1.3 | 3.59e^-40^ |
| 19 | Adrenocortical carcinoma (77) | GTEx healthy adrenal gland tissue (127) | -1.3 | 1.14e^-13^ |
| 20 | Kidney papillary cell carcinoma (288) | Normal adjacent tissue (32) | -1 | 1.79e^-21^ |
| **Comparisons with no significant changes in *DMD* gene expression** | | | | |
| 21 | Kidney chromophobe cell carcinoma (66) | Normal adjacent tissue (25) | -0.2 | 1 |
| 22 | Acute myeloid leukemia (173) | GTEx healthy whole blood (337) | 0.04 | 1 |
| 23 | Kidney clear cell carcinoma (530) | Normal adjacent tissue (72) | 0.08 | 1 |
| **Comparisons with increased expression of the *DMD* gene in tumor samples** | | | | |
| 24 | Thyroid carcinoma (504) | Normal adjacent tissue (59) | 0.9 | 1.89e^-14^ |
| 25 | Diffuse large B-cell lymphoma (47) | GTEx healthy whole blood (337) | 4.8 | 2.40e^-22^ |

Supplementary Table 2. Number of tumor and control tissues in clusters generated by cutting the dendrogram of the hierarchical clustering analysis at different levels. Six is highlighted as it was chosen to be the optimal number of clusters.

| Number of clusters at different dendrogram cut-off levels | Tissue type | Composition of clusters | | | | | | |
| --- | --- | --- | --- | --- | --- | --- | --- | --- |
|  |  | 1 | 2 | 3 | 4 | 5 | 6 | 7 |
| 3 | Tumor tissues | 10 | 1 | 16 |  |  |  |  |
|  | Control tissues | 2 | 0 | 20 |  |  |  |  |
| 4 | Tumor tissues | 10 | 1 | 14 | 0 |  |  |  |
|  | Control tissues | 2 | 0 | 20 | 2 |  |  |  |
| 5 | Tumor tissues | 9 | 1 | 14 | 1 | 0 |  |  |
|  | Control tissues | 2 | 0 | 20 | 0 | 2 |  |  |
| 6 | Tumor tissues | 9 | 1 | 12 | 1 | 2 | 0 |  |
|  | Control tissues | 2 | 0 | 4 | 0 | 16 | 2 |  |
| 7 | Tumor tissues | 7 | 1 | 12 | 2 | 1 | 2 | 0 |
|  | Control tissues | 1 | 0 | 4 | 1 | 0 | 16 | 2 |

Supplementary Table 3. *DMD* gene expression levels in 18 healthy tissues from the GTEx database represented as a percentage of two housekeeping genes (HKGs): *PGK1* and *HMBS*. The percentage of *DMD* level of each HKG in healthy tissues divided by the percentage of *DMD* of each HKG in skeletal muscle is also displayed.

| Healthy tissue  (Number of samples) | | *PGK1* | | *HMBS* | |
| --- | --- | --- | --- | --- | --- |
|  |  | *DMD*% of PGK1 | *DMD*% of *PGK1* in tissue / *DMD*% of *PGK1* in muscle | *DMD*% of *HMBS* | *DMD*% of *HMBS* in tissue / *DMD*% of *HMBS* in muscle |
| 1 | Skeletal Muscle (396) | 94.6 | 100.0 | 138.2 | 100.0 |
| 2 | Pancreas (165) | 94.6 | 100.0 | 122.1 | 88.3 |
| 3 | Uterus (78) | 93.3 | 98.6 | 134.7 | 97.4 |
| 4 | Colon (304) | 89.2 | 94.3 | 127.3 | 92.1 |
| 5 | Bladder (9) | 88.0 | 93.0 | 131.6 | 95.2 |
| 6 | Ovary (88) | 85.7 | 90.5 | 137.4 | 99.4 |
| 7 | Stomach (173) | 84.8 | 89.6 | 120.7 | 87.3 |
| 8 | Breast (179) | 84.5 | 89.3 | 125.3 | 90.6 |
| 9 | Liver (110) | 84.5 | 89.3 | 111.5 | 80.7 |
| 10 | Prostate (100) | 84.0 | 88.8 | 117.1 | 84.7 |
| 11 | Esophagus (651) | 79.1 | 83.6 | 121.5 | 87.9 |
| 12 | Skin (557) | 78.3 | 82.8 | 117.0 | 84.6 |
| 13 | Thyroid (278) | 77.7 | 82.2 | 110.1 | 79.6 |
| 14 | Cervix (10) | 77.5 | 81.9 | 115.0 | 83.2 |
| 15 | Adrenal Glands (127) | 76.4 | 80.8 | 108.5 | 78.5 |
| 16 | Lung (287) | 73.8 | 78.0 | 111.4 | 80.6 |
| 17 | Kidney (27) | 65.2 | 68.9 | 95.5 | 69.1 |
| 18 | Whole Blood (337) | 30.8 | 32.5 | 43.7 | 31.6 |

Supplementary Table 4. *DMD* gene expression in 13 TCGA primary tumors compared to NATs from the TCGA and healthy GTEx tissues. P-values were calculated using the UCSC Xena Browser and adjusted for multiple testing using the Bonferroni correction. P-values were multiplied by the total numbers of tests (n= 13) and compared with the overall level of α= 0.05. Statistically non-significant comparisons are highlighted in grey.

| TCGA tumors  (Number of samples) | | TCGA NATs | | | GTEx healthy tissues | | |
| --- | --- | --- | --- | --- | --- | --- | --- |
|  |  | Number of samples | LogFC | *p*-value | Number of samples | LogFC | *p*-value |
| 1 | BRCA (1,092) | 113 | -3.7 | 7.66e^-115^ | 179 | -2.5 | 5.23e^-173^ |
| 2 | BLCA (407) | 19 | -3.5 | 1.26e^-05^ | 9 | -4.0 | 2.53e^-04^ |
| 3 | UCEC (180) | 23 | -3.1 | 7.65e^-15^ | 78 | -3.7 | 1.61e^-64^ |
| 4 | COAD (288) | 41 | -1.8 | 5.64e^-11^ | 304 | -3.2 | 6.45e^-96^ |
| 5 | STAD (414) | 36 | -1.7 | 1.78e^-04^ | 173 | -1.5 | 1.17e^-37^ |
| 6 | PRAD (495) | 52 | -1.6 | 2.23e^-14^ | 100 | -0.9 | 3.33e^-14^ |
| 7 | LUSC (498) | 50 | -1.6 | 3.94e^-48^ | 287 | -1.3 | 5.77e^-53^ |
| 8 | LIHC (369) | 50 | -1.5 | 9.32e^-37^ | 110 | -0.9 | 4.62e^-13^ |
| 9 | LUAD (513) | 59 | -1.3 | 1.86e^-40^ | 287 | -1.0 | 4.39e^-47^ |
| 10 | KIRP (288) | 32 | -1 | 9.30e^-22^ | 27 | -0.6 | 1 |
| 11 | KICH (66) | 25 | -0.2 | 1 | 27 | 0.3 | 0.7 |
| 12 | KIRC (530) | 72 | 0.08 | 1 | 27 | 0.9 | 2.77e^-08^ |
| 13 | THCA (504) | 59 | 0.9 | 9.84e^-15^ | 278 | 0.5 | 6.84e^-11^ |

Supplementary Table 5. Expression of four *DMD* gene transcripts in 25 TCGA primary tumors and corresponding NATs from the TCGA or healthy tissues form the GTEx database. P-values were calculated using the UCSC Xena Browser using a two-tailed Welch’s t test and adjusted for multiple testing using the Bonferroni correction. P-values were multiplied by the total numbers of tests (n= 25) and compared with the overall level of α= 0.05. Statistically non-significant comparisons are highlighted in grey.

| Pairwise comparisons (Number of samples) | | | Dp427m | | Dp71 | | Dp71b | | Dp71ab | |
| --- | --- | --- | --- | --- | --- | --- | --- | --- | --- | --- |
|  |  |  | LogFC | *p*-value | LogFC | *p*-value | LogFC | *p*-value | LogFC | *p*-value |
| 1 | Uterine carcinosarcoma (57) | GTEx normal uterus tissue (78) | -7.2 | 1.29e^-18^ | 0.2 | 1 | -0.6 | 0.1 | 1 | 0.3 |
| 2 | Bladder urothelial carcinoma (407) | Normal adjacent tissue (19) | -7.1 | 3.31e^-07^ | 0.2 | 7.11e^-06^ | -3 | 9.29e^-06^ | -0.04 | 1 |
| 3 | Breast invasive carcinoma (1,092) | Normal adjacent tissue (113) | -7 | 7.97e^-149^ | 0.4 | 1 | -2.7 | 3.58e^-84^ | -1.9 | 7.60e^-06^ |
| 4 | Lung squamous cell carcinoma (498) | Normal adjacent tissue (50) | -6.6 | 6.29e^-32^ | -0.7 | 1 | -2 | 3.03e^-37^ | -1.8 | 1.57e^-06^ |
| 5 | Cervical and endocervical cancer (303) | GTEx healthy cervix tissue (10) | -6.5 | 3.87e^-3^ | 0.3 | 1 | -1.2 | 5.83e^-3^ | 0.009 | 1 |
| 6 | Uterine corpus endometrioid carcinoma (180) | Normal adjacent tissue (23) | -6.3 | 7.94e^-09^ | 0.2 | 3.83e^-2^ | -2.9 | 4.26e^-16^ | 1.2 | 2.78e^-14^ |
| 7 | Head and neck squamous cell carcinoma (518) | Normal adjacent tissue (44) | -5.2 | 1.11e^-08^ | 1.2 | 1.86e^-2^ | -0.9 | 1.02e^-3^ | 0.1 | 1 |
| 8 | Adrenocortical carcinoma (77) | GTEx healthy adrenal gland tissue (125) | -4.8 | 2.09e^-20^ | 0.03 | 1 | -0.7 | 3.09e^-2^ | -2.1 | 7.63e^-08^ |
| 9 | Ovarian serous cystadenocarcinoma (419) | GTEx healthy ovary tissue (88) | -4.6 | 6.75e^-45^ | 0.3 | 1 | -1.3 | 1.36e^-31^ | -0.9 | 2.97e^-2^ |
| 10 | Colon adenocarcinoma (287) | Normal adjacent tissue (41) | -4.4 | 1.58e^-07^ | 0.4 | 2.98e^-08^ | -0.9 | 8.75e^-07^ | -0.4 | 1 |
| 11 | Kidney papillary cell carcinoma (288) | Normal adjacent tissue (32) | -4 | 2.71e^-06^ | -0.5 | 1 | -1.1 | 1.59e^-14^ | -3.4 | 2.19e^-07^ |
| 12 | Esophageal carcinoma (181) | GTEx healthy esophagus tissue (650) | -3.9 | 1.87e^-22^ | 2.4 | 4.22e^-23^ | -0.6 | 1.84e^-05^ | 4 | 2.35e^-50^ |
| 13 | Stomach adenocarcinoma (413) | Normal adjacent tissue (36) | -3.8 | 1.71e^-05^ | -1.1 | 1 | -0.4 | 1 | 0.3 | 1 |
| 14 | Kidney chromophobe cell carcinoma (66) | Normal adjacent tissue (25) | -3.5 | 9.66e^-4^ | -0.3 | 1 | -0.3 | 1 | -0.6 | 1 |
| 15 | Lung adenocarcinoma (513) | Normal adjacent tissue (59) | -3.4 | 2.35e^-09^ | 0.2 | 1 | -1.2 | 3.78e^-24^ | -1 | 4.05e^-02^ |
| 16 | Prostate adenocarcinoma (494) | Normal adjacent tissue (51) | -3.2 | 9.40e^-10^ | -0.4 | 1 | -1.6 | 5.27e^-15^ | -3.2 | 2.18e^-06^ |
| 17 | Thyroid carcinoma (504) | Normal adjacent tissue (59) | -2.7 | 4.64e^-07^ | 0.2 | 1 | 1.1 | 3.91e^-13^ | -0.4 | 1 |
| 18 | Rectal adenocarcinoma (92) | Normal adjacent tissue (10) | -5.9 | 0.1 | 0.4 | 8.40e^-3^ | -1 | 0.1 | 0.4 | 1 |
| 19 | Kidney clear cell carcinoma (530) | Normal adjacent tissue (72) | -1.1 | 0.4 | 0.9 | 4.13e^-2^ | -0.1 | 1 | -1.2 | 1.57e^-4^ |
| 20 | Liver hepatocellular carcinoma (369) | Normal adjacent tissue (50) | -0.6 | 1 | 0.1 | 1 | -1.2 | 6.76e^-18^ | -0.1 | 1 |
| 21 | Acute myeloid leukemia (173) | GTEx healthy whole blood (336) | -0.04 | 1 | 0.8 | 3.09e^-08^ | 1.3 | 2.70e^-11^ | 0.7 | 4.82e^-07^ |
| 22 | Skin cutaneous melanoma (102) | GTEx healthy skin tissue (556) | 0.1 | 1 | -0.3 | 1 | -1.2 | 3.49e^-10^ | -0.2 | 1 |
| 23 | Cholangiocarcinoma (36) | Normal adjacent tissue (9) | 1.3 | 1 | -0.1 | 1 | -2.3 | 7.48e^-08^ | 0.9 | 1 |
| 24 | Diffuse large B-cell lymphoma (47) | GTEx healthy whole blood (336) | 0.7 | 1 | 1.5 | 2.43e^-3^ | 6.8 | 6.14e^-22^ | 3.2 | 2.12e^-09^ |
| 25 | Pancreatic adenocarcinoma (178) | GTEx healthy pancreas tissue (165) | 3.6 | 3.39e^-22^ | -0.9 | 2.13e^-10^ | -2 | 6.53e^-48^ | -2.4 | 3.17e^-23^ |

Supplementary Table 6. Expression of Dp71 and its splice variants in primary blood malignancies in the TARGET database compared to GTEx healthy whole blood. P-values were calculated using the UCSC Xena Browser using a two-tailed Welch’s t test and adjusted for multiple testing using the Bonferroni correction. P-values were multiplied by the total numbers of tests (n= 2) and compared with the overall level of α= 0.05.

| Pairwise comparisons (Number of samples) | | | Dp427m | | Dp71 | | Dp71b | | Dp71ab | |
| --- | --- | --- | --- | --- | --- | --- | --- | --- | --- | --- |
|  |  |  | LogFC | *p*-value | LogFC | *p*-value | LogFC | *p*-value | LogFC | *p*-value |
| 1 | Acute lymphoblastic leukaemia (37) | GTEx healthy whole blood (336) | 1.9 | 3.83e^-04^ | 2.4 | 1.71e^-04^ | 0.7 | 2.59e^-02^ | 1.9 | 3.83e^-04^ |
| 2 | Acute myeloid leukaemia (29) | GTEx healthy whole blood (336) | 2.3 | 1.67e^-05^ | 2.6 | 7.68e^-07^ | 0.9 | 3.74e^-02^ | 2.3 | 1.67e^-05^ |

### Supplementary Table 7. Values of Pearson’s correlation between the expression of dystrophin protein and other members of the DAP complex (those encoded by *SNTB2*, *DAG1*, *DTNBP1*, *DTNA*, and *SNTA1*). Correlation values were calculated using the Data Explorer tool at the DepMap portal across all tumor cell lines using data from the “Proteomics” database.

| Protein encoded by | Pearson’s coefficient | Q-value | Number of cell lines |
| --- | --- | --- | --- |
| *SNTB2* | 0.527 | 33.05e^-24^ | 366 |
| *DAG1* | 0.397 | 4.14e^-12^ | 366 |
| *DTNBP1* | -0.321 | 1.56e^-08^ | 366 |
| *DTNA* | 0.368 | 2.60e^-12^ | 348 |
| *SNTA1* | 0.380 | 7.57e^-09^ | 216 |

Supplementary Table 8. Differentially expressed miRNAs between samples with low vs. high *DMD* gene expression that were common in five or more tumors out of 15 different primary tumor types (FDR-corrected p-value < 0.1, Bonferroni-corrected p-value < 0.05, |LogFC| value >= 0.5).

| Downregulated miRNAs | Number of tumors (out of 15) | Upregulated miRNAs | Number of tumors (out of 15) |
| --- | --- | --- | --- |
| hsa-miR-1247-3p | 7 | hsa-miR-33a-5p | 6 |
| hsa-miR-143-3p | 7 | hsa-miR-96-5p | 6 |
| hsa-let-7c-3p | 6 | hsa-miR-141-3p | 5 |
| hsa-miR-125b-5p | 6 | hsa-miR-15a-5p | 5 |
| hsa-miR-139-3p | 6 | hsa-miR-17-5p | 5 |
| hsa-miR-145-5p | 6 | hsa-miR-19a-3p | 5 |
| hsa-miR-217 | 6 | hsa-miR-200a-3p | 5 |
| hsa-miR-30a-3p | 6 |  |  |
| hsa-miR-143-5p | 5 |  |  |
| hsa-miR-381-3p | 5 |  |  |
| hsa-miR-99a-5p | 5 |  |  |
